# Lewy-MSA hybrid fold drives distinct neuronal α-synuclein pathology

**DOI:** 10.1101/2024.11.21.624748

**Authors:** Masahiro Enomoto, Ivan Martinez-Valbuena, Shelley L. Forrest, Xiaoxiao Xu, Renato P. Munhoz, Jun Li, Ekaterina Rogaeva, Anthony E. Lang, Gabor G. Kovacs

**Affiliations:** Princess Margaret Cancer Centre, University Health Network, Toronto, ON, M5G 1L7, Canada; Tanz Centre for Research in Neurodegenerative Diseases, University of Toronto, Toronto, Canada; Department of Medical Biophysics, University of Toronto, Toronto, Canada; Edmond J. Safra Program in Parkinson’s Disease and the Morton and Gloria Shulman Movement Disorders Clinic, Toronto Western Hospital, M5T 2S8, Toronto, Ontario, Canada; Krembil Brain Institute, University Health Network, Toronto, M5T 0S8, Ontario, Canada; Department of Laboratory Medicine and Pathobiology, University of Toronto, Toronto, Canada; Laboratory Medicine Program, University Health Network, Toronto, Canada

## Abstract

The ordered assembly of α-synuclein protein into filaments encoded by SNCA characterizes neurodegenerative diseases called synucleinopathies. Lewy body disease (LBD) shows predominantly neuronal α-synuclein pathology and multiple system atrophy (MSA) predominantly oligodendrocytic α-synuclein pathology affecting subcortical brain structures. Based on cryo-electron microscopy, it was reported that structures of α-synuclein filaments from LBD differ from MSA and juvenile onset synucleinopathy (JOS) caused by a 21-nucleotide duplication in the second exon of one allele of *SNCA* gene ^1-3^. Importantly, a rare subtype of MSA, called atypical MSA^4^ shows abundant neuronal argyrophilic α-synuclein inclusions in the limbic system. Current concepts indicate that disease entities are characterized by unique protofilament folds. Here we demonstrate that in addition to the MSA fold, α-synuclein can form a new Lewy-MSA hybrid fold in the same brain region, leading to the atypical histopathological form of MSA. Distinct biochemical characteristics of α-synuclein, as demonstrated by protease-sensitivity digestion assay, seed amplification assays (SAAs) and conformational stability assay (CSA), are also linked to cytopathological differences (e.g. neuronal or oligodendroglial). We expand the current structure-based classification of α−synucleinopathies and propose that cell-specific protein pathologies can be associated with distinct filament folds.

## INTRODUCTION

Description of argyrophilic oligodendroglial cytoplasmic inclusions called Papp-Lantos bodies linked three syndromes, striatonigral degeneration, olivopontocerebellar atrophy and Shy-Drager syndrome, as various manifestations of the same disease, MSA^5^. The filamentous inclusions contain α-synuclein^6^ as seen also in LBD. Later, clinical types of MSA with parkinsonism (MSA-P) or predominant cerebellar ataxia (MSA-C) were defined^7^. In addition to distinct solubility in SDS of α-synuclein filaments of MSA from those of LBD^8^, differences in seeding capacity^9^ and the protofilament folds have been described^1,3^.

Neuronal cytoplasmic inclusions (NCIs) can be also found in MSA, mostly in the brainstem and subcortical structures^10^ and are thought to contribute to memory impairment when seen in the hippocampus^11^. Following reports of a temporal variant of MSA with large argyrophilic neuronal inclusions^12-15^, the concept of atypical MSA or frontotemporal lobar degeneration (FTLD)-synuclein was proposed^4^. Some studies suggested that frequent globular neuronal cytoplasmic inclusions in the medial temporal region are important pathological finding of MSA cases with dementia grouped as those with and without distinct FTLD^16^. Patients with Pick body-like α-synuclein immunoreactive inclusions show more severe atrophy of the medial temporal lobes and broader spreading of NCIs than those without^17^.

## RESULTS

Based on neuropathology we identified the peculiar histopathological features of atypical MSA and evaluated the clinical files. The clinical diagnosis was compatible with MSA-P. Importantly, cognitive decline was not evident clinically, although detailed neuropsychology testing was not performed. Brain MRI performed 14 months prior to the death showed bilateral putaminal atrophy more left than right (**Extended data suppl-Fig. 1**). Additionally, prominent asymmetric atrophy was noted in the mesial temporal lobe, predominantly in the amygdala region. Genetic analysis using Neurobooster array did not detect any known pathogenic mutations in genes linked to neurodegenerative diseases, including multiplications at the *SNCA* locus.

Neuropathology examination (see also **Extended data suppl-Fig. 1**) showed round and horseshoe-like NCIs visible in Hematoxylin and eosin staining in the hippocampus CA1 (**Fig. 1a**), subiculum, granule cell layer of the dentate gyrus (**Fig. 1b**), and amygdala (**Fig. 1c, d**) and in neocortical areas. NCIs (cortex, hippocampus, amygdala, striatum, brainstem), neuronal nuclear inclusions (putamen), and typical Papp-Lantos bodies in the cortical, subcortical and cerebellar white matter and putamen were strongly immunoreactive for disease-associated α-synuclein (**Fig. 1 e-o**). Asymmetric involvement was noted, including the amygdala (**Fig. 1c, d and g, h**) and frontal cortex (**Fig. 1i, j**). In summary, α-synuclein pathology was compatible with MSA. Although a striatonigral degeneration pattern was recognized, the prominent neuronal α-synuclein pathology involving the limbic system and the severe atrophy was striking. Based on the morphology and distribution of neuronal α-synuclein pathology^18^, atypical MSA was diagnosed. In addition, neurofibrillary degeneration was compatible with Braak stage II without Aβ positive plaques interpreted as primary-age related tauopathy (PART)^19^. The presence of 4-repeat tau-immunopositive argyrophilic grains and oligodendroglial coiled-bodies in the limbic white matter was compatible with argyrophilic grain disease (Saito stage II)^20^. There was a lack of phosphorylated TDP-43 pathology, Aβ plaques and cerebral amyloid angiopathy.

**Figure 1.**
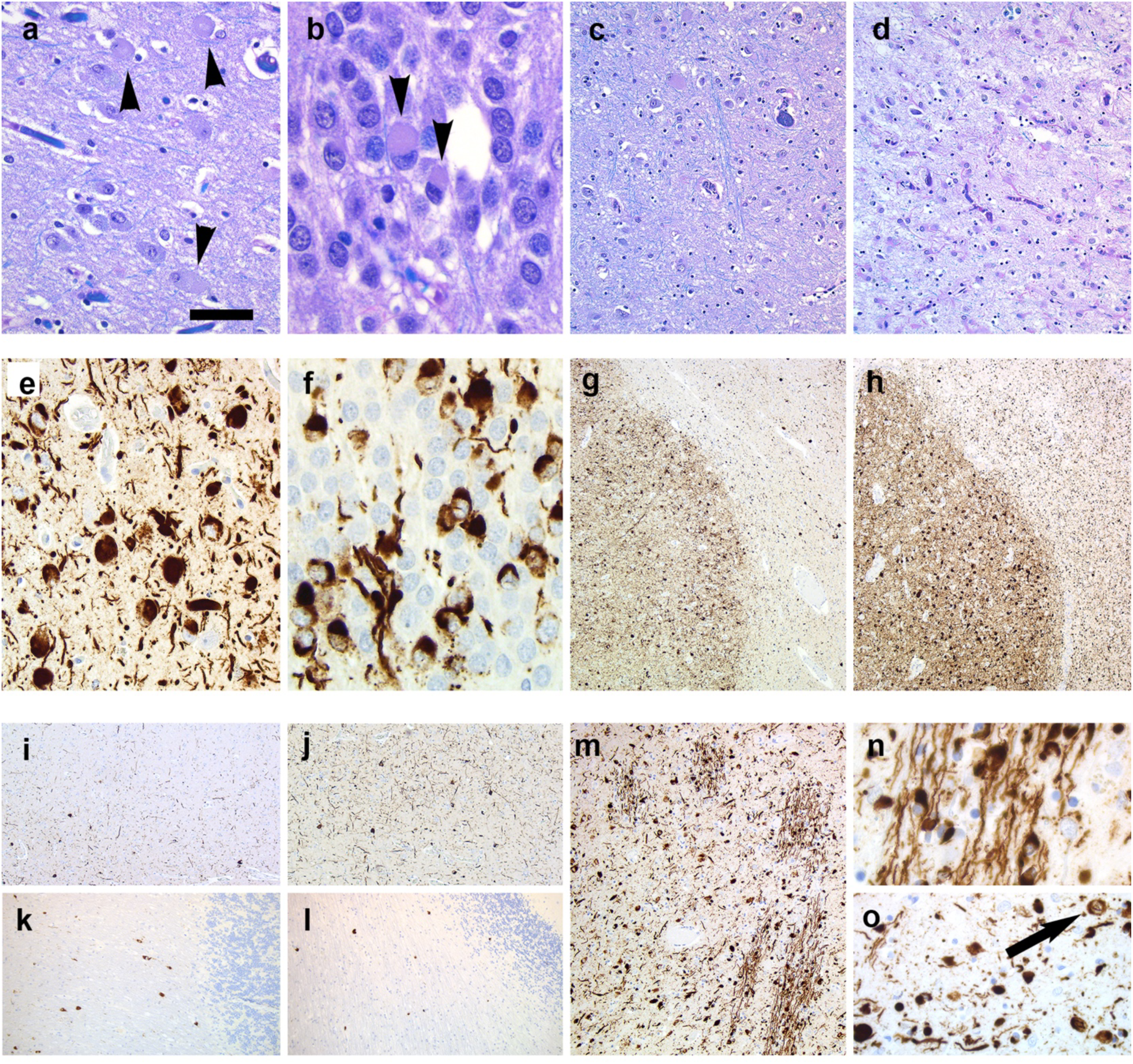
Neuropathological findings. Round and horseshoe-like NCIs (indicated by black arrowheads) visible in Hematoxylin and eosin staining (**a, b**) and immunostaining for disease-associated α-synuclein (**e, f**) in the hippocampus CA1 (**a**), granule cell layer of the dentate gyrus (**b**). Asymmetric involvement of the amygdala showing neuronal cytoplasmic inclusions and ballooned neurons in the left side (**c**) and severe gliosis in the right side (**d**) corresponding to differences in the load of α-synuclein pathology in the left (**g**) and right (**h**). Immunostaining for disease-associated α-synuclein reveals less neuronal cytoplasmic inclusions and thread-like pathology in the left (**i**) than right (**j**) frontal cortex and more Papp-Lantos bodies in the left (**k**) than right (**l**) cerebellar white matter. The putamen shows severe α-synuclein pathology (**m-o**) in the form of enlarged Papp-Lantos bodies (**n**), neuronal cytoplasmic and nuclear inclusions (**o**; arrow). Bar in figure (**a**) represents 40 µm for a, e, o; 20 µm for b, f; 80 µm for c, d; and m, 25 µm for n; 200 µm for g, h, i, j; and 120 µm for k and l.

After identifying the unique neuronal α-synuclein inclusions, we next sought to assess whether this was associated with distinct biochemical, physical, and seeding properties compared to those already described for synucleinopathies. For these analyses, samples from the cerebellar white matter and temporal cortex of the atypical MSA patient, as well as cerebellar white matter from an individual showing the classical form of MSA and temporal cortex from an LBD patient (**Extended data suppl-Table 1**) were dissected and processed into sarkosyl-insoluble and detergent-soluble homogenates. Immunostaining for α-synuclein in these four regions, performed on tissue from the same hemisphere as that used for frozen tissue extraction, showed abundant Papp-Lantos bodies in the cerebellar white matter of the classical form of MSA and less in the case with atypical MSA (**Fig. 2a, b**), neuronal inclusions in the temporal cortex of the atypical MSA patient (**Fig. 2c**), and Lewy bodies and neurites in the temporal cortex of the LBD patient (**Fig. 2d**). Our strategy of collecting samples for histology and biochemistry from the same side was crucial since there was significant asymmetry in the involvement of limbic structures as we reported earlier for other diseases as well^21^. We then examined biochemical, physical, and seeding differences in the α-synuclein present across these synucleinopathies. Using immunoblotting under denaturing conditions, we assessed the sarkosyl-insoluble extracts for total α-synuclein (**Fig. 2e**). In all four samples α-synuclein monomers (∼14 kDa) were present; however, high molecular weight α-synuclein species as well as lower molecular weight bands were uniquely detected in the sarkosyl-insoluble extract from the atypical MSA temporal cortex (**Fig. 2e**). To further explore the physical properties of α-synuclein in these sarkosyl-insoluble extracts, we performed a protease-sensitivity digestion assay with 40 μg/ml proteinase K, a method previously used to define different α-synuclein conformations^22^. Post-proteinase K digestion, α-synuclein patterns diverged significantly. While the sarkosyl-insoluble α-synuclein from the cerebellum of both MSA and atypical MSA cases was fully digested, a portion of the sarkosyl-insoluble α-synuclein in the temporal cortex of the LBD case resisted digestion (**Fig. 2f**). Notably, the majority of sarkosyl-insoluble α-synuclein from the temporal cortex of atypical MSA was proteinase K-resistant (**Fig. 2f**). Next, we assessed whether differences in α-synuclein seeding capacity existed between these four samples. Using seed amplification assays (SAAs) that are able to quantify the propensity of misfolded α synuclein for aggregation, tracking increases in the thioflavin T (ThT) fluorescence at multiple time points^9^. We measured five kinetic parameters from each reaction: lag time, growth phase, T50 (time to reach 50% of maximum aggregation), ThT max, and the area under the curve (AUC) of the fluorescence response. Our findings showed similar seeding capacities in the cerebellar samples from both MSA cases, while α-synuclein seeds from the LBD temporal cortex exhibited a slower T50. Notably, α-synuclein seeds from the atypical MSA temporal cortex demonstrated the fastest aggregation rate and highest ThT max, suggesting a distinct seeding capacity of the α-synuclein present in this region (**Fig. 2g**).

**Figure 2.**
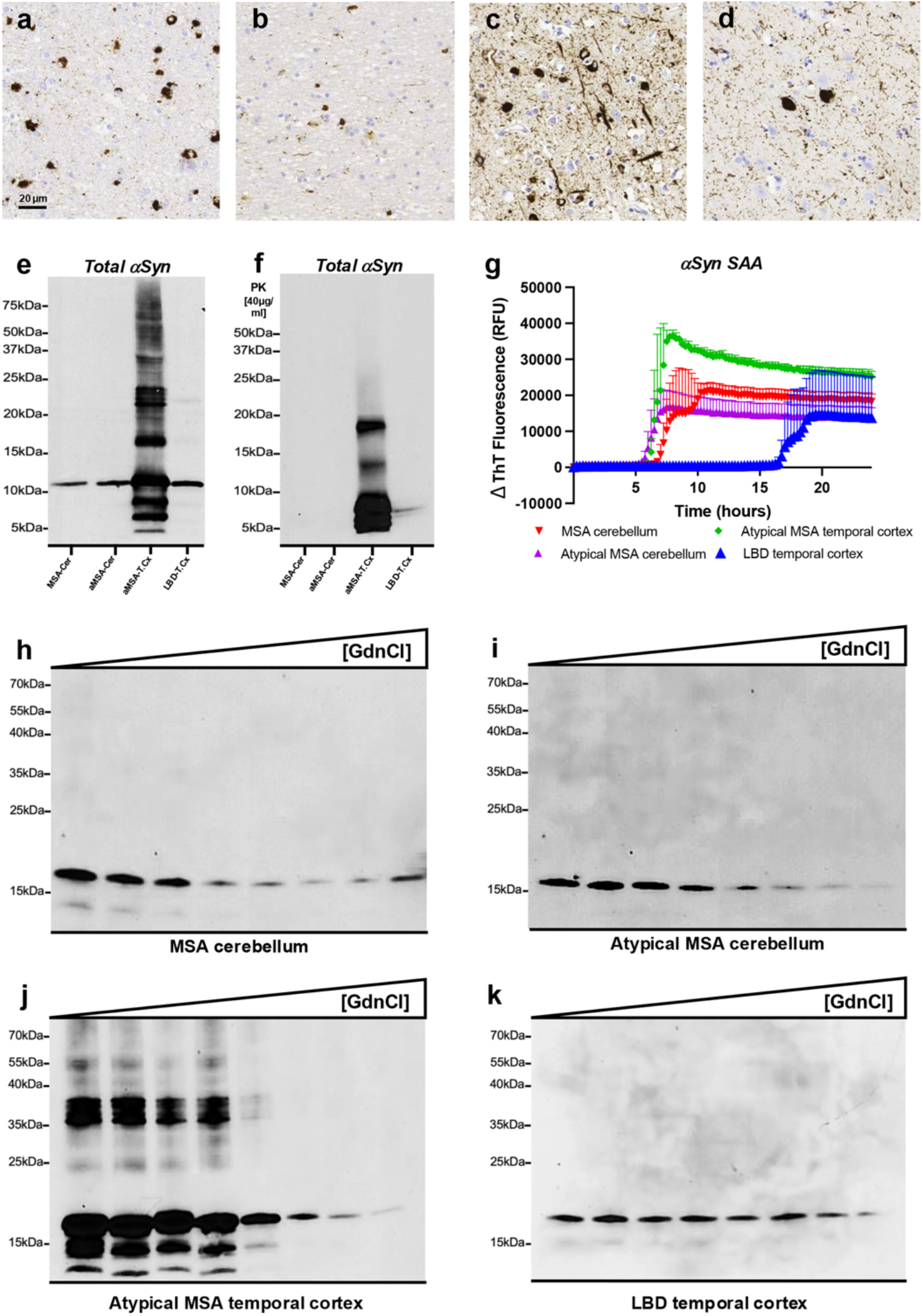
Biochemical Characterization of α-Synuclein in the Cerebellum and Temporal Cortex of atypical MSA. Representative immunohistochemistry images for aggregated α-synuclein are shown in the cerebellar white matter of an MSA patient displaying GCIs (**a**), in the cerebellar white matter of the atypical MSA patient showing GCIs (**b**), in the temporal cortex of the atypical MSA patient with neuronal inclusions (**c**), and in the temporal cortex of an LBD patient with Lewy bodies and neurites (**d**). An immunoblot for total α-synuclein (Syn-1 clone) illustrates the distinct banding pattern of the sarkosyl-insoluble filaments (**e**), along with the results of proteinase K (PK) digestion of the same sarkosyl-insoluble filaments isolated from the cerebellum of the MSA and atypical MSA patients and the temporal cortex of the atypical MSA and LBD patients, respectively (**f**). Aggregation curves of α-synuclein in the presence of sarkosyl-insoluble filaments from the same samples used in the immunoblot are provided, with data shown as the mean from representative subjects measured in quadruplicate (**g**). Conformational stability assay (CSA) results for α-synuclein aggregates in detergent-soluble brain extracts are depicted for the same patients, with representative α-synuclein immunoblots and the resultant denaturation curves shown for the cerebellum of the MSA patient (**h**), the cerebellum of the atypical MSA patient (**i**), the temporal cortex of the atypical MSA patient (**j**), and the temporal cortex of the LBD patient (**k**). Cer: cerebellum; T. Cx.: temporal cortex; SAA: seed amplification assay; GdnCl: guanidine hydrochloride.

Finally, using conformational stability assay (CSA), which measure protein aggregate strain stability in the presence of guanidine hydrochloride (GdnCl), we observed that α-synuclein aggregates from the classical form of MSA and atypical MSA cerebellar extracts were significantly less stable than those from the LBD temporal cortex (**Fig. 2h, i, k**), consistent with previous findings^9^. However, CSA on α-synuclein aggregates from the atypical MSA temporal cortex revealed a unique banding pattern. In samples treated with 1, 1.5, 2, and 2.5 M GdnCl, we consistently detected a ∼17 kDa α-synuclein band alongside several lower molecular weight bands and GdnCl-resistant α-synuclein aggregates between 35 and 40 kDa (**Fig. 2j**). Together, these data suggest that α-synuclein in the temporal cortex of atypical MSA may be structurally distinct from that in the cerebellum of the same patient and in the classical neuropathology form of MSA and LBD cases.

To confirm this hypothesis, we next evaluated filament types derived from the temporal cortex of the atypical MSA case using cryogenic electron microscopy (cryo-EM). The majority of α-synuclein filaments isolated from this region comprised a doublet filament, which has a left-handed helical twist (**Extended Data suppl-Fig. 1**). The structure of these filaments was determined at 3.2 Å resolution (**Extended Data suppl-Fig. 2**). The cryo-EM map and the local resolution estimate showed that there are two separate major regions in one rung of the filament, the higher- and the lower-resolution region (**Fig. 3a, b**). Model building revealed two distinct protofilaments that asymmetrically pack against each other (**Fig. 3c**). One protofilament is composed of the Lewy core fold and the MSA fold^1,3^ termed as “Lewy-MSA hybrid protofilament” in which the Lewy core fold is the part of the higher-resolution region and the MSA fold covers almost the entire lower-resolution region in **Fig. 3b**. To the best of our knowledge, the Lewy-MSA hybrid is the first structure that comprises two distinct folds identified from seemingly different diseases (e.g. LBD and MSA) in a single protofilament. The other protofilament also holds MSA fold (hereafter referred to as the MSA protofilament), which covers the rest of the higher-resolution region (**Fig. 3c**). Although the estimated local resolution of the lower-resolution region is 3.3-4.0 Å (**Fig. 3b**), we succeeded in building a model of the MSA fold of the Lewy-MSA hybrid protofilament with the 0.51 of Cα model-to-map correlation. The residues of structures modeled in cryo-EM map are shown in **Fig. 4a**. In the typical Lewy fold the C-terminus (residues E61-L100) packs with the Lewy core fold (residues G31-K60) (**Fig. 4c**). Contrarily, the Lewy core (residues G25-E46) of the Lewy-MSA hybrid protofilament is packed with the MSA protofilament (**Fig. 4b**), which makes the MSA fold of the Lewy-MSA hybrid protofilament isolated from the Lewy core flexible, causing a lower resolution structure. The C-terminus (residues E61-L100) in the typical Lewy fold interacts with the Lewy core fold (residues G31-K60)^3^ (**Fig. 4c**). As described previously, two major subtypes are known in the MSA fold, MSA Type I and MSA Type II^1^. Both subtypes comprise two non-identical protofilaments named PF-IA, PF-IB, PF-IIA and PF-IIB, respectively. The MSA fold of the MSA protofilament is PF-IIB, as it includes the K43-E46 short β-strand found in PF-IIB but not in PF-IB among known MSA folds (**Fig. 4d**). The MSA fold of the Lewy-MSA hybrid protofilament could be also categorized as PF-IIB because it has the identical short β-strand of K43-E46 that is shared as a part of the Lewy core (**Fig. 4c**). Hereafter, for the sake of simplicity, the Lewy and MSA folds in the Lewy-MSA hybrid protofilament are referred to as Lewy-L and Lewy-M, respectively as shown in **Fig. 4b**. In summary, the Lewy-MSA hybrid protofilament is composed of Lewy-L and Lewy-M folds. The MSA fold in the other protofilament is tightly packed with Lewy-L (**Fig. 4b**).

**Figure 3.**
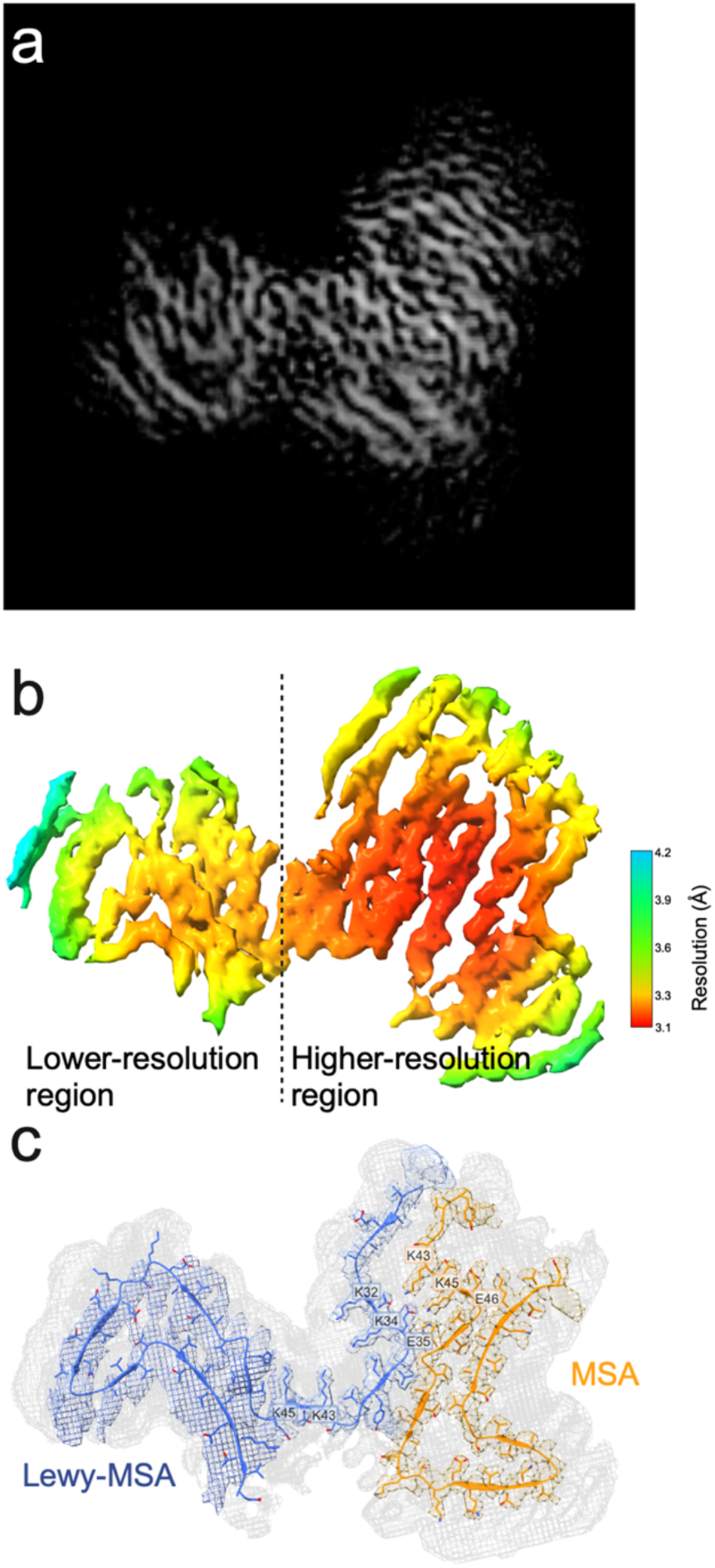
Cryo-EM maps and atomic model of Lewy-MSA hybrid α-synuclein filament. (**a**) The cross-section of the cryo-EM map of Lewy-MSA hybrid α-synuclein filament from the temporal cortex of the atypical MSA patient. (**b**) The local resolution estimation of cryo-EM map of Lewy-MSA hybrid filament. (**c**) Sharpened high-resolution cryo-EM maps of Lewy-MSA hybrid α-synuclein filament with overlaid the atomic model. Unsharpened, 4.5 Å low pass-filtered map is in grey. The sharpened maps with different counter levels are used for Lewy-L/MSA complex folds and Lewy-MSA fold. The boundary is G47 of the Lewy-MSA protofilament.

**Figure 4.**
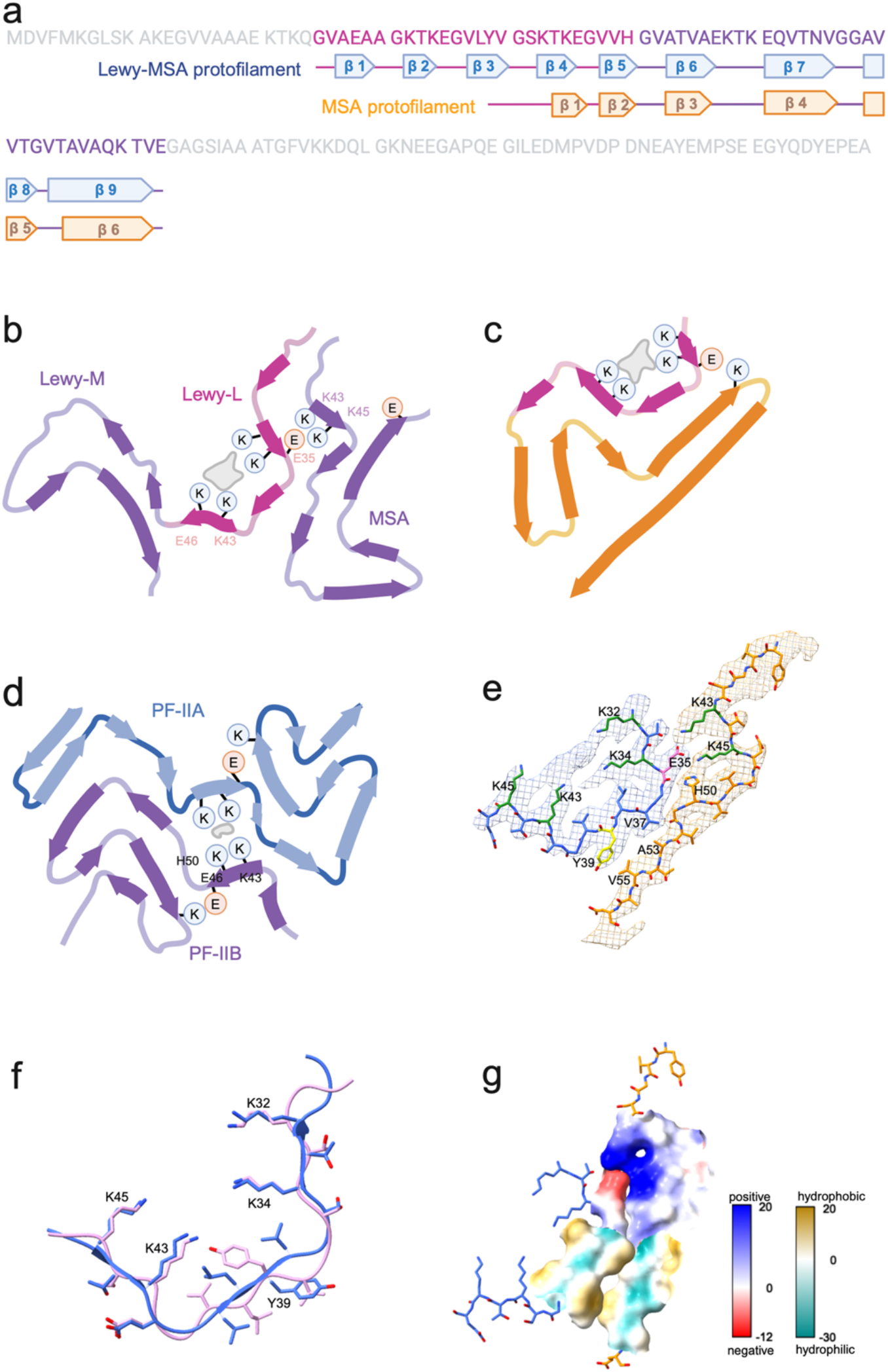
Cryo-EM structure of Lewy-MSA hybrid α-synuclein filament. (**a**) Sequence of human α-synuclein (in grey). The residues of which structures were modeled in cryo-EM map are shown in pink/purple. N-terminal region (residues 25-46) of the Lewy-MSA hybrid protofilament corresponding to Lewy core (pink) shows similarity to Lewy body and the C-terminal region (residues 47-83) in purple shows similarity to MSA-PF-IIB. The secondary structure of Lewy-MSA hybrid protofilament is shown in arrow boxes (blue for Lewy-MSA and orange for MSA). (**b, c, d**) Comparison of the Lewy-MSA hybrid fold with the Lewy and MSA-II α-synuclein filament folds. Schematic of secondary structure elements in the Lewy-MSA hybrid (**b**), the Lewy (**c**) and MSA-II (**d**) folds, depicted as a single rug. The Lewy-MSA hybrid fold (**b**), the Lewy core (**c**) and MSA-PF-IIB (**d**) folds are coloured as in (**a**). Thick connecting lines with arrowheads indicate β-strands. (**e**) Close-up view of the inter-protofilament interface of Lewy-L and MSA and Lewy-L around a non-proteinaceous strong density. In the interface between Lewy-L and MSA, E35 (magenta) and Y39 (yellow) of Lewy-L and K43 and K45 (green) of MSA are highlighted. In Lewy core of Lewy-L K32, K34, K43 and K45 are highlighted in green. (**f**) Electrostatic potential and hydrophobicity maps of the inter-protofilament interface of Lewy-L and MSA.

As expected from the results shown in **Fig. 2**, we have determined a totally new structure, the Lewy-MSA/MSA doublet filaments (**Fig. 4b**) that is distinct from those identified previously from MSA and LBD cases (**Fig. 4c, d**). The key question about this new filament structure is how does it exhibit a distinct proteinase K-resistance and protein aggregate stability (**Fig. 2**)? The difference in the inter-protofilament interface of the two protofilaments could be an answer. The two non-identical protofilaments of MSA Type I and Type II folds pack against each other through an extended interface that forms a large cavity surrounded by the side chains of K43, K45 and H50 from each protofilament (**Fig. 4d, e**). This cavity encloses a non-proteinaceous strong density, the chemical nature of which remains to be established^1^. As opposed to the known α-synuclein doublet filament structure from human brains^1,3^, the Lewy-MSA hybrid fold has a closer interface between two protofilaments (**Fig. 4e, f**). There is no additional strong density in the interface between Lewy-L and MSA folds. Instead, K43 and K45 of the MSA fold directly interact with E35 of Lewy-L through the electrostatic interaction. The CH-π interaction between K45 and H50 of MSA could secondly reinforce this electrostatic interaction^1^. Further, the hydrophobic interactions among V37 and Y39 of the Lewy-L and A53 and V55 of MSA maintain this interface closer. This stable interface is connected to the rest of the MSA mainly through the electrostatic interaction between E46 and K80. As far as we are aware, the Lewy-MSA/MSA doublet filaments structure identified in this study is the first example from human brains which does not contain any non-proteinaceous density in the inter-protofilament interface. As opposed to the interface between Lewy-L and MSA, we confirmed that the cavity of the Lewy-L encloses a non-proteinaceous strong density surrounded by the side chains of K32, K34, K43 and K45 (**Fig. 3c** and **4b, d**) as described in the typical Lewy fold^1^. The root-mean-square deviation (RMSD) value of all the atoms (excluding hydrogens) of K32-K45 except V37-V40 between Lewy-L and the typical Lewy core is 1.133, which indicates the Lewy-L is almost identical with the typical Lewy core except for Y39 that faces towards the inter-protofilament interface and forms hydrophobic interaction with V55 of the MSA (**Fig. 4e-g**). Collectively, we found the novel doublet α-synuclein filament, Lewy-MSA hybrid fold that exhibits a distinctive structure in which the Lewy core (Lewy-L) is crammed with two MSA-PF-IIB folds (Lewy-M and MSA) (**Fig. 4a-d**).

Previous studies reported that α-synuclein filaments adopt the same structures in different individuals with MSA, LBD and JOS^1-3^ similar to tau filaments from the brains of individuals with various tauopathies^23-27^. Altogether, this suggested that neurodegenerative disease-related protein filaments adopt an identical fold in individuals with the same disease, but different subtypes of the same proteinopathies are characterised by distinct folds^1,27^. Our results indicate that distinct folds including hybrid forms (e.g. MSA fold and Lewy-MSA hybrid fold) can be present in the same brain and that the predominant cytopathologies can be associated with distinct folds. The eosinophilic round Pick-body-like inclusions also resemble cortical Lewy bodies^15^ but show strong Gallyas-Braak silver staining positivity. Previously, the predominance of Type I and II filaments of MSA fold were found in different brain regions, e.g. putamen and cerebellum, respectively, and it was not concluded whether MSA Type I and Type II filaments are common to both nerve cells and glial cells^1^.

Here we expand the folds described in MSA and show that hybrid folds consisting of protofilaments of two seemingly different disease (e.g. MSA and LBD) can exist in the same brain. This could suggest that the predominance of cytopathologies (e.g. neuronal or oligodendroglial) is associated with distinct folds of neurodegenerative disease-related proteins that can be identified also based on distinct biochemical characteristics and seeding capacity. Although the temporal cortex of the atypical MSA case in this study exhibits the presence of the MSA fold, the typical MSA doublet filaments were not identified. This could also suggest that the cofactors in the Lewy core fold and in the inter-protofilament interface of the MSA Type I and Type II might be different, implying that the cofactors could be molecular determinants of α-synuclein filaments structures which drive distinct human α-synuclein pathologies. It will be important to explore whether this phenomenon is present in other neurodegenerative proteinopathies.

## Methods

### Data reporting

No statistical methods were used to predetermine sample size. The experiments were not randomized, and the investigators were not blinded to allocation during experiments and outcome assessment.

### Clinical history

A 68-year-old right-handed person was assessed for potential advanced therapies at the Movement Disorders Clinic of the Toronto Western Hospital. He had a four-year history of parkinsonism, first noticed in the form of global limitations in mobility and voice changes. He was initially diagnosed as Parkinson Disease (PD) and started on levodopa. Treatment, however, was never felt to be beneficial overall and his general condition deteriorated progressively. Importantly, certain symptoms became important sources of disability, including dysarthria and hypophonia, dysphagia causing choking episodes, and postural instability with frequent falls leading to two hospital admissions. Over time, levodopa dose was increased, added by entacapone and rotigotine, with modest, if any, benefit. He denied having clear motor fluctuations, dyskinesias or dystonia. He did not recognize changes in sense of smell or taste, sleep was fragmented with symptoms of sleep apnea, for which CPAP was indicated. No signs of REM-sleep behavior disorder (RBD) were identified. There were no subjective cognitive abnormalities, no psychosis or behavioral changes. For the past two to three years, he experienced erectile disfunction, episodes of bladder and bowel incontinence, and symptomatic orthostatic hypotension. Past medical history was positive for longstanding anxiety and depression, coronary artery disease, hypertension and hypothyroidism. Treatment regimen included levodopa 1250 mg a day divided in 5 intakes, entacapone 200 mg with all doses of levodopa, rotigotine patch 8mg a day, escitalopram 10 mg a day, dutasteride 0.5 mg a day, tamsulosin 0.4 mg a day, ASA 81 mg a day, atorvastatin 40 mg a day, bisoprolol 5 mg twice a day, mirtazapine 45 mg at bedtime. On examination, the patient was alert and oriented, speech was significantly hypophonic and dysarthric, difficult to understand. Saccades were slow with mild limitation in vertical smooth pursuit. Remaining cranial nerve exam was unremarkable. Muscle bulk was globally reduced, no fasciculations were observed. There were bilateral, mild postural, fine, jerky distal movements (“polyminimyoclonus”); muscle tone was mildly increased symmetrically in all extremities, moderately increased in the neck; there was moderate symmetric bradykinesia. Deep tendon reflexes were symmetrical, and brisk throughout but no pathological hyperreflexia. Plantar reflexes were flexor bilaterally. Coordination and sensory exams were normal. He was brought to the clinic in a wheelchair, required assistance to stand but was able to stand with support of a cane or walker for short distance ambulation. MoCA test scored 27/30 (2 months before death); however, detailed neuropsychology evaluation was not performed. The duration of illness was 5 years.

### Genetic analysis

Genomic DNA was isolated from brain using a QIAGEN kit. The entire open reading frame of *SNCA* was Sanger sequenced (**Extended data and suppl-Table 2**). Gene dosage of the *SNCA* locus on chromosome 4 was determined by inspecting the B allele frequency and LogR Ratio plots (**Extended data suppl-Fig. 3**) from Neurobooster array data (155 variants) using GenomeStudio (Illumina Inc., CA).

### Neuropathology

Brain tissue was examined in the frame of an ongoing Brain Donation program of the Toronto Western Hospital following the approval of the local ethics committee (University Health Network Research Ethics Board, Nr. 20-5258) and adherence to appropriate consent protocols.

Formalin-fixed, 4.5 μm paraffin-embedded tissue sections comprising the frontal, temporal, parietal, anterior cingulate, motor and occipital cortices, anterior and posterior regions of the basal ganglia, thalamus, midbrain, pons, medulla oblongata and cerebellum with dentate nucleus were stained with hematoxylin, eosin, and luxol fast blue (HE/LFB), and Gallyas Braak silver staining. For the neuropathology analysis including the evaluation for mixed pathology, various immunostains including p-tau (clone AT8, 1:1000, Invitrogen/ThermoFisher, Carlsbad, USA), Aβ (Clone 6F/3D, 1:50, Dako, Glostrup, Denmark), a-synuclein (clone 5G4, 1:4000, Analytikjena, Jena, Germany)^28^, phosphorylated TDP-43 (clone 11-9, 1:2000, CosmoBio, Tokyo, Japan) and anti-p62 (3/P62 LCK LIGAND, 1:500 BD Company, NJ, United States) were employed. Alzheimer’s Disease (AD)-related pathology was assessed through the Braak neurofibrillary tangle (NFT) stage^29^ and Thal phase for Aβ deposition^30^. To evaluate any Lewy body disease-related lesions we adhered to consensus criteria using α-synuclein immunohistochemistry^31^. AGD was classified based on Saito stage^20^. Detailed neuropathology report is provided in **Extended data suppl-Fig.1**.

### Protein extraction

Detergent-extracted brain homogenates [10% (w/v)] were prepared in PBS using a gentle-MACS Octo Dissociator (Miltenyi BioTec). Nine volumes of brain homogenate were then mixed with one volume of 10× detergent buffer [5% (v/v) Nonidet P-40, 5% (w/v) sodium deoxycholate in PBS] containing Pierce Universal Nuclease (ThermoFisher) and Halt Phosphatase Inhibitor (ThermoFisher), and then incubated on ice for 20 min. Samples were clarified by centrifugation at 1000x *g* for 5 min at 4 °C to generate detergent-extracted brain homogenate, as previously described.^22^ The supernatant was collected and aliquoted in 0.5-ml low protein binding tubes (Eppendorf) to avoid excessive freeze–thaw cycles. A bicinchoninic acid protein (BCA) assay (ThermoFisher) was performed to determine the total protein concentration of all samples.

The sarkosyl-insoluble extraction was performed as previously described, with minor modifications^1^. Briefly, 1 gram of tissue was homogenised in 20% (v/w) extraction buffer consisting of 10 mM Tris-HCl, pH 7.5, 0.8 M NaCl, 10% sucrose and 1 mM EGTA (Sigma). Homogenates were brought to 2% sarkosyl (Sigma) and incubated for 30 min. at 37 °C. Following a 13 min. centrifugation at 10,000 g, the supernatants were spun at 100,000 g for 30 min. The pellets were resuspended in 500 μl/g extraction buffer and centrifuged at 3,000 g for 5 min. The supernatants were diluted 3-fold in 50 mM Tris-HCl, pH 7.5, containing 0.15 M NaCl, 10% sucrose and 0.2% sarkosyl (Sigma), and spun at 166,000 g for 40 min. Sarkosyl-insoluble pellets were resuspended in 100 μl/g of PBS (Invitrogen).

### α-Synuclein seed amplification assay (α-Syn SAA)

SAA reactions were performed in 384-well plates with a clear bottom (Nunc) as previously described.^9^ Briefly, full-length human recombinant α-Synuclein (Impact Biologicals) was thawed from −80 ºC storage, reconstituted in HPLC-grade water (Sigma) and filtered through a 100-kDa spin filter (ThermoFisher) in 500 μl increments. 10 μl of the biological sample (containing 0.005 μl of the sarkosyl-insoluble fraction dissolved in H_2_0) was added to the wells containing 20 μl of the reaction buffer (0.1 M PB, pH 8, 0.875 M Na_3_Citrate), 10 μl of 50 μM ThT and 10 μl of 0.5 mg/ml of monomeric recombinant α-Syn. The plate was sealed and incubated at 37 °C in a BMG FLUOstar Omega plate reader with cycles of 1 min shaking (400 rpm double orbital) and 14 min rest. ThT fluorescence measurements (450±10 nm excitation and 480±10 nm emission, bottom read) were taken every 15 min for a period of 30 h. Each sample was tested in quadruplicate.

### SDS-PAGE and immunoblotting

Gel electrophoresis was performed using 4-12% or 12% Bolt Bis-Tris Plus gels (Thermo Scientific). Proteins were transferred to 0.45-μm nitrocellulose membranes for 60 min at 30 V. Proteins were crosslinked to the membrane via 4% (v/v) paraformaldehyde incubation in PBS for 30 min at room temperature, with rocking. The membranes were blocked for 60 min at room temperature in blocking buffer (5% [w/v] skim milk in 1× TBST (TBS and 0.05% [v/v] Tween-20)) and then incubated overnight at 4 °C with primary antibodies directed against amino acids 15–123 of the α-Synuclein protein (1:1,000 dilution, ref: 610786, BD Biosciences) diluted in the blocking buffer. The membranes were washed three times with TBST and then incubated for 60 min at room temperature with horseradish peroxidase-conjugated secondary antibodies (1:3,000 dilution, ref: 172-1011, Bio-Rad) in the blocking buffer. Following another three washes with TBST, immunoblots were developed using Western Lightning enhanced chemiluminescence Pro (PerkinElmer) and imaged using X-ray films.

### Proteinase K digestions

Protease digestions were performed as previously described^9,22^ with minor modifications. A concentration of 40 μg/ml of proteinase K were added to the sarkosyl-insoluble homogenates. Samples were incubated at 37 °C with continuous shaking (600 rpm) for 60 min. Proteinase K digestions were halted with a final concentration of 4 mM PMSF. Samples were resuspended in 1× LDS buffer (Life Technologies) and analyzed by SDS-PAGE followed by immunoblotting, as described above.

### Conformational stability assay (CSA)

Twenty microliters of 2× guanidine hydrochloride (GdnCl) stocks were added to an equal volume of the detergent -extracted brain homogenates to yield final GdnCl concentrations of 0, 1, 1.5, 2, 2.5, 3, 3.5 and 4 M, as previously described^9,22^. Briefly, detergent-extracted brain homogenates were incubated at room temperature with shaking for 120 min (800 rpm) before being diluted to 0.4 M GdnCl in PBS. Following high-speed ultracentrifugation at 100,000 *g* for 60 min at 4 ºC, the pellets were resuspended in 1× LDS loading buffer and boiled for 10 min. Levels of residual α-synuclein were determined by SDS–PAGE followed by immunoblotting, as mentioned above. **Electron cryo-microscopy** Extracted α-synuclein filaments were applied to homemade glow-discharged holey carbon gold grids coated with homemade graphene oxide and plunge-frozen in liquid ethane using Thermo Fisher Vitrobot Mark IV. Micrographs were acquired on a Thermo Fisher Titan Krios microscope operated at 300 kV using Thermo Fisher Falcon 4i direct electron detector in a counting mode. Further details are provided in **Extended data suppl-Table 3**.

### Helical reconstruction

Movie frames were corrected for beam-induced motion and dose-weighted using RELION’s motion-correction implementation^32^. Aligned, non-dose-weighted micrographs were used to estimate the contrast transfer function using CTFFIND-4.1^33^. All subsequent image-processing steps were performed using helical reconstruction methods in RELION^34,35^. α-synuclein filaments were picked using crYOLO Version 1.9.9^36^. Segments with a box size of 1,024 pixels and a pixel size of 1.03 Å comprising an entire helical crossover were extracted using an inter-box distance of 14.1 Å and downscaled to 256 pixels. The segments were initially separated by reference-free 2D classification and those contributing to suboptimal 2D class averages were discarded. An initial helical twist of -1.4° was calculated from the apparent crossover distance of the filaments in 2D class averages and helical rise was fixed at 4.75 Å. An initial 3D reference model was generated *de novo* from 2D class averages of segments that comprise an entire helical crossover using the *relion_helix_inimodel2d* program^34^. The segments were then re-extracted using a box size of 320 pixels without downscaling. The segments were further separated by reference-free 2D classification and those contributing to suboptimal 2D class averages were discarded. The first 3D auto-refinement was carried out using these segments and an initial *de novo* model low-pass filtered to 10 Å. Combination of 3D classification and 3D auto-refinement for several rounds with optimization of T value, helical twist and rise showed separation of β-strands along the helical axis. We then performed Bayesian polishing and contrast transfer function refinement to further improve the resolution of the reconstruction. The final reconstruction was sharpened using standard post-processing procedures in RELION^34^. Overall resolution estimates were calculated from Fourier shell correlations at 0.143 between the two independently refined half-maps, using phase randomisation to correct for convolution effects of a generous, soft-edged solvent mask^37^. Local resolution estimate was obtained using the same phase randomisation procedure. An initial atomic model (PDB-ID 9E9X) was built using Model Angelo^38^ and manually checked and refined in Coot^39^ and ISOLDE^40^. Coordinates refinements were performed using Phenix^41^. The final model was validated with MolProbity^42^. Figures were prepared with ChimeraX^43^. Model statistics are given in **Extended data suppl-Table 3**.

## Data Availability Statement

The cryo-EM map in this study has been deposited in the Electron Microscopy Data Bank (EMDB) with the accession number EMD-47820. The corresponding refined atomic model has been deposited in the Protein Data Bank (PDB) under accession number PDB:9E9X.

## Acknowledgements

We thank the family of the patient for arranging brain donation. Research reported in this publication was supported by the National Institute on Aging of the National Institutes of Health under Award Number R01AG080001, the Edmond J. Safra Foundation, and the Rossy Family Foundation. G.G.K. holds the Rossy chair for PSP research at UHN. SLF is supported by the National Health and Medical Research Council of Australia Ideas grant (#214090508). We thank the Toronto High-Resolution High Throughput cryo-EM facility, supported by the Canada Foundation for Innovation and Ontario Research Fund, for cryo-EM data collection. We also thank Talos L120C funded by Canada Foundation for Innovation (CFI) and Ontario Research Fund-Research Innovation (ORF-RI), for screening cryo-EM samples.

## Author contributions

R.M, A.EL., S.L.F., and G.G.K., identified the patient, collected clinical information (R.M, A.EL.,) and performed neuropathology (S.L.F. and G.G.K.). E.R performed genetic analysis. I.M-V. and J.L. prepared α-synuclein filament samples and performed biochemical analyses. M.E. and X.X. performed cryo-EM data acquisition, performed cryo-EM structure determination. G.G.K. supervised the project. All authors contributed to writing the manuscript.

## Ethics statement

The brain was obtained at autopsy through appropriate consenting procedures with Local Ethical Committee approval. This study was approved by the UHN Research Ethics Board (Nr. 20-5258) and was performed per the ethical standards established in the 1964 Declaration of Helsinki, updated in 2008.

## Additional information

Supplementary information: The online version contains supplementary material.

**Correspondence and requests for materials** should be addressed to Gabor G. Kovacs.

