## Extended data for "Lewy-MSA hybrid fold drives distinct neuronal α-synuclein pathology"

**Suppl-Table 1**. Summary of cases examined for biochemistry and seeding.

| Case | Age | Sex | Aβ Thal phase | NFT Braak stage | CAA: Cerebral amyloid angiopathy | Lewy Body Braak Stage | TDP-43 pathology (LATE-NC) | Other |
|---|---|---|---|---|---|---|---|---|
| **atypical MSA** | 68 | M | - | Stage II | - | - | - | Argyrophilic grain disease |
| **MSA-P** | 64 | F | Phase 5 | Stage II | Aβ-positive Type 1 | - | - | - |
| **LBD** | 60 | M | Phase 1 | Stage III | - | 6 | - | Argyrophilic grain disease |

**Suppl-Table 2. Genomic analysis**

Genomic DNA for the atypical MSA cases was isolated from brain using a QIAGEN kit. The entire open reading frame of the *SNCA* gene was Sanger sequenced using the primers shown the table. Gene dosage of the *SNCA* gene region on chromosome 4 was determined by inspecting the B allele frequency and LogR Ratio plots (Figure) of Neurobooster array data of 155 SNPs in genomestudio (Illumina Inc., CA).
Table. Sequences of primers used in Sanger sequencing of *SNCA*.

| primer name | primer sequence 5'-3' |
|---|---|
| **SNCA_exon1_F** | TTTGTCCAATGGTTAAATGAGTG |
| **SNCA_exon1_R** | CCTTTTTGACAAGCAATGATG |
| **SNCA_exon2_F** | CCTCCTGTTAGCTGGGCTTT |
| **SNCA_exon2_R** | GCTCAGTGATTGTTTTACAATTTCA |
| **SNCA_exon3_F** | TGGCTTTTGTTTCTTCTGACC |
| **SNCA_exon3_R** | GTAGCCGTTCCCCACAGTAA |
| **SNCA_exon4_F** | CGGAGGCATTGTGGAGTTTA |
| **SNCA_exon4_R** | TGCAAGTTGTCCACGTAATGA |
| **SNCA_exon5_F** | GCAGAATATTTGCAAAAACATTGAT |
| **SNCA_exon5_R** | GAAGCACCGAAATGCTGAGT |

**Suppl-Table 3.** Statistics for cryo-EM data acquisition and atomic model building.

| | Lewy-MSA (EMD-47820, PDB 9E9X) |
|---|---|
| **Data acquisition** | |
| Microscope | Titan Krios G3 |
| Detector | Falcon 4i |
| Magnification | 75,000 |
| Voltage (kV) | 300 |
| Exposure rate (e/pix/s) | 6.8 |
| Electron dose (e/Å$^2$) | 41 |
| Defocus range (μm) | 1.3-3.0 |
| Pixel size (Å) | 1.03 |
| **Map refinement** | |
| Symmetry imposed | C1 |
| Initial particle images (No.) | 155,610 |
| Final particle images (No.) | 47,227 |
| Map resolution (Å) | 3.2 |
| FSC threshold | 0.143 |
| Helical twist (°) | -1.2857 |
| Helical rise (Å) | 4.68955 |
| **Model refinement** | |
| Model resolution (Å) | 3.3 |
| FSC threshold | 0.5 |
| Map sharpening *B* factor (Å$^2$) | -50.2323 |
| Model composition | |
| Non-hydrogen atoms | 2,178 |
| Protein residues | 312 |
| Ligands | 0 |
| B factors (Å$^2$) | |
| Protein | 33.55 |
| R.m.s. deviation | |
| Bond lengths (Å) | 0.002 |
| Bond angles (°) | 0.491 |
| Validation | |
| MolProbity score | 2.07 |
| Clashscore | 10.22 |
| Poor totamers (%) | 1.20 |
| Ramachandran plot | |
| Favored (%) | 92.16 |
| Allowed (%) | 7.84 |
| Disallowed (%) | 0 |

### Detailed neuropathology report

Neuropathology examination showed mild loss of neurons in the frontal (suppl-**Suppl-Fig 1f**), motor (**Suppl-Fig 1g**), and temporal cortex. The caudate nucleus was preserved (**Suppl-Fig 1h**), but the putamen showed severe gliosis (**Suppl-Fig 1i**), while the globus pallidus did not show neuronal loss (**Suppl-Fig 1j**). The left-side hippocampus had a normal morphology without mentionable gliosis (**Suppl-Fig 1k**). The right-side hippocampus was not available for histology (only frozen sample). Eosinophilic neuronal cytoplasmic inclusions were detected mostly in the granule cells of the dentate gyrus (**Suppl-Fig 1l**). The amygdala showed severe gliosis and eosinophilic inclusion bodies in neurons (**Suppl-Fig 1m**) with more prominent changes in the right side. Ballooned neurons were noted in the amygdala. The substantia nigra (**Suppl-Fig 1n**) and pontine base showed gliosis and patchy Purkinje cell loss was noted in the cerebellum (**Suppl-Fig 1o**). Brainstem type Lewy bodies were not seen. Immunostaining for α-synuclein showed many neuronal cytoplasmic inclusions and threads of the frontal (**Suppl-Fig 1p**) and temporal cortex. This was associated with a lower number of Papp-Lantos bodies in the white matter (**Suppl-Fig 1q**) of the frontal and temporal but more in the motor area. Prominent α-synuclein immunoreactive Papp-Lantos bodies were seen in the internal capsule (**Suppl-Fig 1r**), subinsular peri-putamen white matter and putamen, and less in the cerebral peduncles, pontine base, medulla oblongata, and the cerebellar white matter (**Suppl-Fig 1s**). Frequent cytoplasmic and nuclear neuronal inclusions and threads were noted in the putamen and less in brainstem nuclei. The amygdala exhibited many large spherical neuronal α-synuclein inclusions (**Suppl-Fig 1t**). Immunostaining for α-synuclein, p62 and AT8 also showed prominent pathologies in the hippocampus (**Suppl-Fig 1u-w**). Many α-synuclein positive neuronal inclusions were seen in the CA1 subregion and low amounts of Papp-Lantos bodies were seen in the hippocampal white matter (**Suppl-Fig 1x, y**). Round, Pick-body-like, and ring-like neuronal cytoplasmic α-synuclein positive inclusions predominated in granule cells of the dentate gyrus (**Suppl-Fig 1z1**); many of these showed p62-immunoreactivity (**Suppl-Fig 1z2**) and were Gallyas-Braak silver positive. In addition, a few granule cells showed fine granular AT8- (**Suppl-Fig 1z3**) but not phosphorylated TDP-43-immunoreactivity (**Suppl-Fig 1z4**). This was accompanied by oligodendrocytic coiled bodies in the hippocampus and amygdala white matter, granular/fuzzy astrocytes in the amygdala, and argyrophilic 4-repeat tau- and p62-immunopositive dendritic grains. Moderate numbers of neurofibrillary tangles and neuropil threads were observed in the entorhinal cortex.

**Suppl-Fig.1. Brain MRI and histopathology (*next page*).**

FLAIR (**a, b**) and T2 weighted (**c-e**) images showing bilateral putamen atrophy and hyperintensity surrounding the putamen (**a-d**) and medial temporal lobe and amygdala atrophy (**e**; indicated by white arrow; right > left). Hematoxylin and eosin and Luxol staining of the frontal (**f**), motor (**g**) cortex, caudate nucleus (**h**), putamen (**i**), globus pallidus (**j**), hippocampus (**k**), dentate gyrus (**l;** arrows indicate eosinophilic inclusions), amygdala (**m**), substantia nigra (**n**), and cerebellum (**o**). Immunostaining α-synuclein (**p-u, x, y,** $\mathbf{z_1}$), p62 (**v,** $\mathbf{z_2}$), phosphorylated tau (**w,** $\mathbf{z_3}$) and phosphorylated TDP-43 ($\mathbf{z_4}$) of the frontal cortex (**p**) and white matter (**q**), internal capsule (**r**), cerebellum white matter (**s**), amygdala (**t**), hippocampus (**u-w**), CA1 subregion (**x**), hippocampal white matter (**y**), and granule cell layer of the dentate gyrus ($\mathbf{z_{1\text{-}4}}$). Bar in (**e**) represents 250 μm for **f-j**, **m**, **n**, **r-t**; 150 μm for **o**, **x**; 100 μm for **p**, **q**; 500 μm for **k**, **u**, **v**, **w**; 50 μm for **l**; 200 μm for **y**; 35 μm for $\mathbf{z_{1\text{-}4}}$.

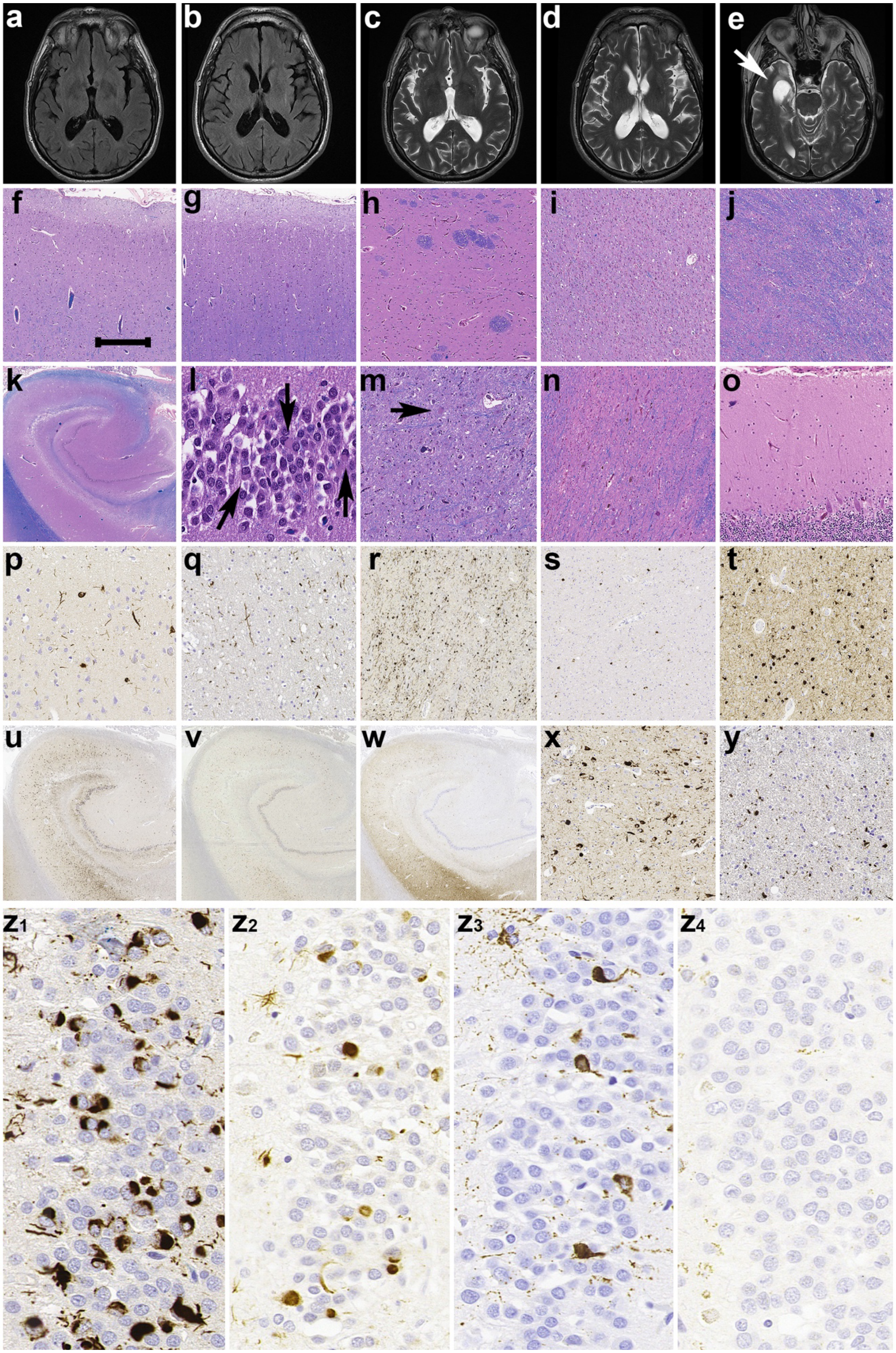
a
b
c
d
e
f
g
h
i
j
k
l
m
n
o
p
q
r
s
t
u
v
w
x
y
z1
z2
z3
z4

**Suppl-Fig.2. Cryo-EM images, maps and resolution evaluation of the refined model.**
(a) A representative cryo-EM image of α-synuclein filament (yellow arrows) from the temporal cortex of an atypical MSA patient. Scale bar, 50 nm. (b) Two-dimensional class average spanning an entire crossover of the Lewy-MSA hybrid filament. Scale bar, 10 nm. (c) Side view of the Lewy-MSA hybrid filament. Scale bar, 2 nm. (d) Fourier shell correlation (FSC) curve of two independently refined half-maps (black line); FSC curve of final cryo-EM reconstruction and refined atomic model (red); FSC curve of first half-map and the atomic model refined against this map (blue); FSC curve of second half-map and the atomic model refined against the first half-map (yellow dashes).

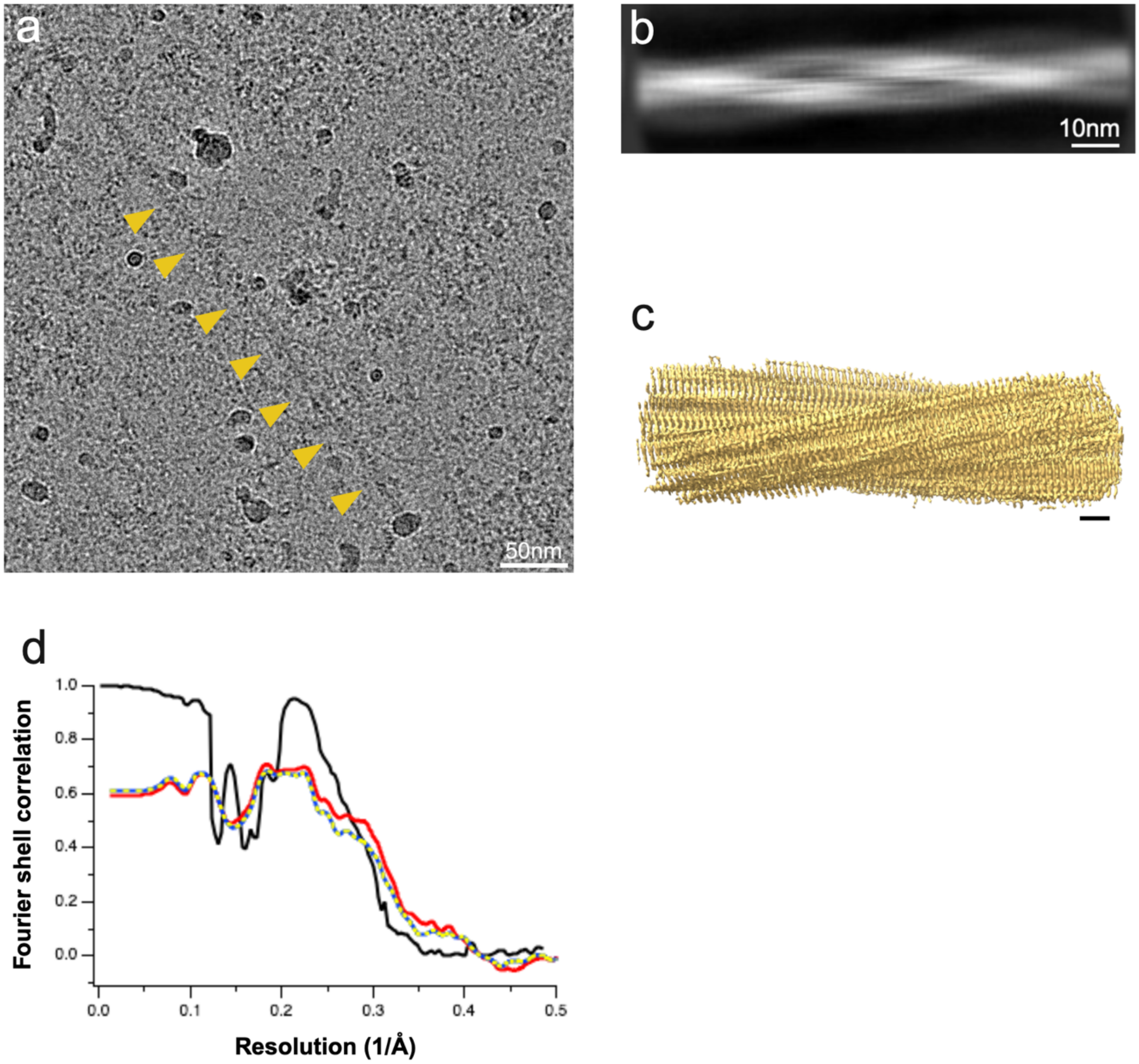

**Suppl-Fig. 3**. Screenshot of B allele frequency and LogR ratio plots for the case with atypical MSA in chromosome browser, genomestudio.

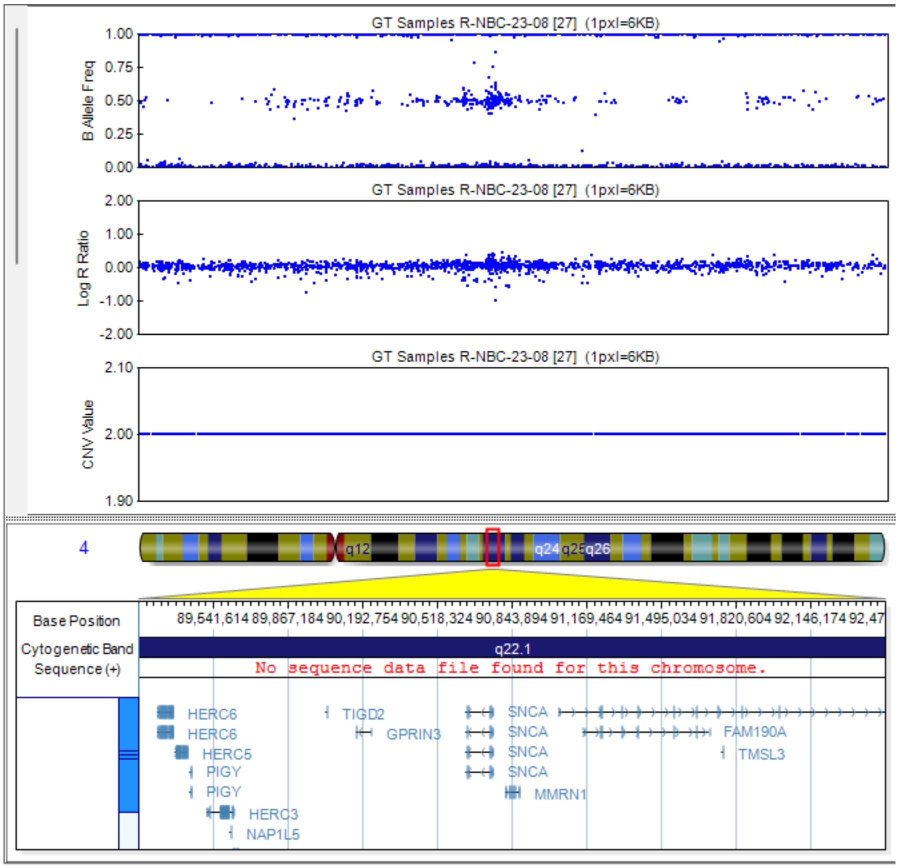
